# Loss of RpoS results in attenuated *Escherichia coli* colonization of human intestinal organoids and a competitive disadvantage within the germ-free mouse intestine

**DOI:** 10.1101/2020.07.30.230003

**Authors:** Madeline R. Barron, Roberto J. Cieza, David R. Hill, Sha Huang, Veda K. Yadagiri, Jason R. Spence, Vincent B. Young

## Abstract

Pluripotent stem-cell-derived human intestinal organoids (HIOs) are three-dimensional, multicellular structures that model a previously uncolonized, naïve intestinal epithelium in an *in vitro* system. We recently demonstrated that microinjection of the non-pathogenic *Escherichia coli* strain, ECOR2, into HIOs induced morphological and functional maturation of the HIO epithelium, including increased secretion of mucins and cationic antimicrobial peptides. In the current work, we use ECOR2 as a biological probe to investigate the bacterial response to colonization of the HIO lumen. In *E. coli* and other Gram-negative bacteria, adaptation to environmental stress is regulated by the general stress response sigma factor, RpoS. We generated an isogenic ∆*rpoS* ECOR2 mutant to compare challenges faced by a bacterium during colonization of the HIO lumen relative to the germ-free mouse intestine, which is currently the best available system for studying the initial establishment of bacterial populations within the gut. We demonstrate that loss of RpoS significantly decreases the ability of ECOR2 to colonize HIOs, though it does not prevent colonization of germ-free mice. Rather, the ∆*rpoS* ECOR2 exhibits a fitness defect in the germ-free mouse intestine only in the context of microbial competition. These results indicate that HIOs pose a differentially restrictive luminal environment to *E. coli* during colonization, thus increasing our understanding of the HIO model system as it pertains to studying the establishment of intestinal host-microbe symbioses.

**Importance:** Technological advancements have and will continue to drive the adoption of organoid-based systems for investigating host-microbe interactions within the human intestinal ecosystem. Using *E. coli* deficient in the RpoS-mediated general stress response, we demonstrate that the type or severity of microbial stressors within the HIO lumen differ from those of the *in vivo* environment of the germ-free mouse gut. This study provides important insight into the nature of the HIO microenvironment from a microbiological standpoint.

## Introduction

The mammalian gastrointestinal tract is inhabited by a diverse community of microbes that play critical roles in host development and health, including facilitating the development and maturation of the intestinal epithelial barrier (1–3). The intestinal epithelium represents an important physical and biochemical interface through which host-microbe symbioses within the gut are established and maintained (4). Historically the model systems available to study intestinal epithelial-microbe interactions consisted primarily of immortalized cell lines and animal models. For instance, germ-free mice have become the gold-standard for investigating host and microbial responses during the establishment of defined bacterial populations within the microbe-naïve gut (5–10). However, technological advancements have expanded the repertoire of systems available for studying host-microbe interactions at the intestinal interface. In this regard, stem-cell-derived human intestinal organoids (HIOs) have emerged as powerful tools to investigate epithelial structure and function following initial interactions with a range of bacterial species (11–13).

HIOs are three-dimensional, organotypic structures comprised of multiple types of differentiated epithelial cells surrounded by a supporting mesenchyme (13–16). Derived from embryonic or induced pluripotent stem cells, HIOs possess many aspects of the microbe-naïve intestinal epithelium in an experimentally tractable, *in vitro* system (13, 15, 16). We recently reported that microinjection of ECOR2, a non-pathogenic strain of *E. coli* originally isolated from a healthy individual (17), into the lumen of HIOs induced transcriptional, morphological, and functional changes in HIOs resulting in maturation of the epithelium (14). These changes included increased expression of genes regulating epithelial tight junction proteins, increased production of mucins, and secretion of antimicrobial peptides (14). Of note, *E. coli* established a stable population within the HIO lumen for several days after microinjection, with preservation of HIO epithelial barrier integrity (14). Overall, this study highlighted the utility of HIOs for studying epithelial adaptation to the establishment of bacterial populations at the intestinal epithelial interface.

Most studies using HIOs have focused on epithelial responses, whereas less attention has been paid to microbial responses modulating colonization of this model of host-microbe symbioses. In the current work, we used *E. coli* as a biological probe to investigate the bacterial response to colonization of the HIO lumen. *E. coli* adapts to its environment via the well-characterized “general stress response”, which is regulated by the stress response sigma factor, RpoS (18–20). Upon exposure to environmental stressors, including nutrient limitation, oxidative stress, and exposure to low pH (19), RpoS alters global bacterial gene expression to switch the cell from a state of active growth to one of survival (19). Here, we generated an isogenic ∆*rpoS* mutant of *E. coli* strain ECOR2 to compare the challenges faced by a bacterium when colonizing the HIO lumen and the murine gut. We demonstrate that loss of RpoS attenuates the ability of ECOR2 to colonize HIOs, though it does not prevent colonization of germ-free mice. Rather, the ∆*rpoS* mutant exhibits a fitness defect in the mouse gut only in the context of microbial competition. Thus, relative to the *in vivo* environment of the germ-free mouse intestine, the HIO lumen is differentially restrictive to *E. coli* during colonization. These results increase our understanding of the HIO model system as it pertains to studying intestinal epithelial-microbe interactions.

## Materials and Methods

### Bacterial strains and plasmids

Strains and plasmids are listed in Table 1. *Escherichia coli* strain ECOR2 (ATCC 35321) (17) was used in the present study. Bacteria were grown aerobically in LB broth or on LB agar at 37°C. When required, growth medium was supplemented with kanamycin (50 ug/mL). An isogenic ∆*rpoS* mutant was constructed by disruption of the ECOR2 *rpoS* gene via lambda red-mediated gene replacement with the aminoglycoside phosphotransferase gene (*neo*) (21) . The *neo* gene was amplified from the plasmid pKD4 (21) using primers MB001-F and MB001-R. The *neo* PCR product containing 50 bases upstream and downstream homologous to the *rpoS* gene was subsequently electroporated into to Red+Gam-producing ECOR2 containing plasmid pKD46 (21). Successful disruption of the *rpoS* gene was verified by end-point PCR using primers MB002-F and MB002-R, Sanger sequencing with primers MB003-F and MB003-R, and real-time quantitative reverse transcription PCR (see below FigureS1A-B in supplemental material).

**Table 1.**
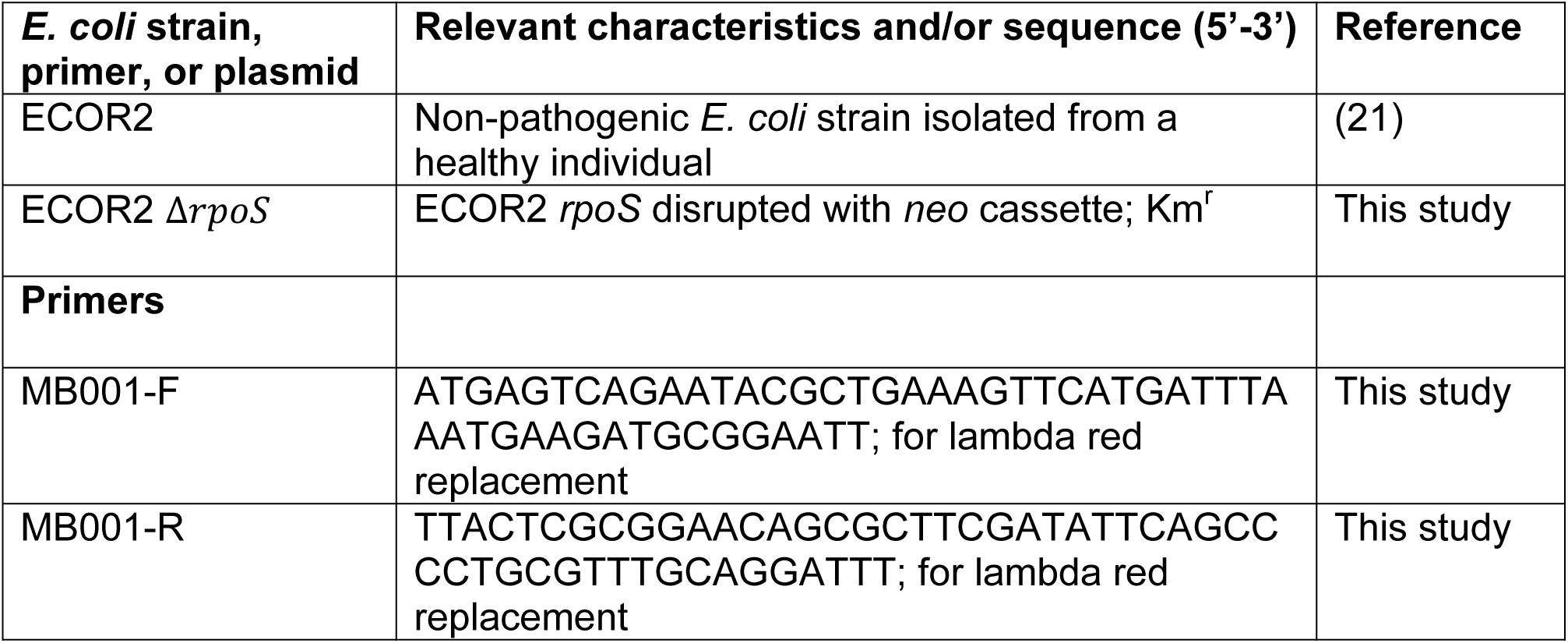
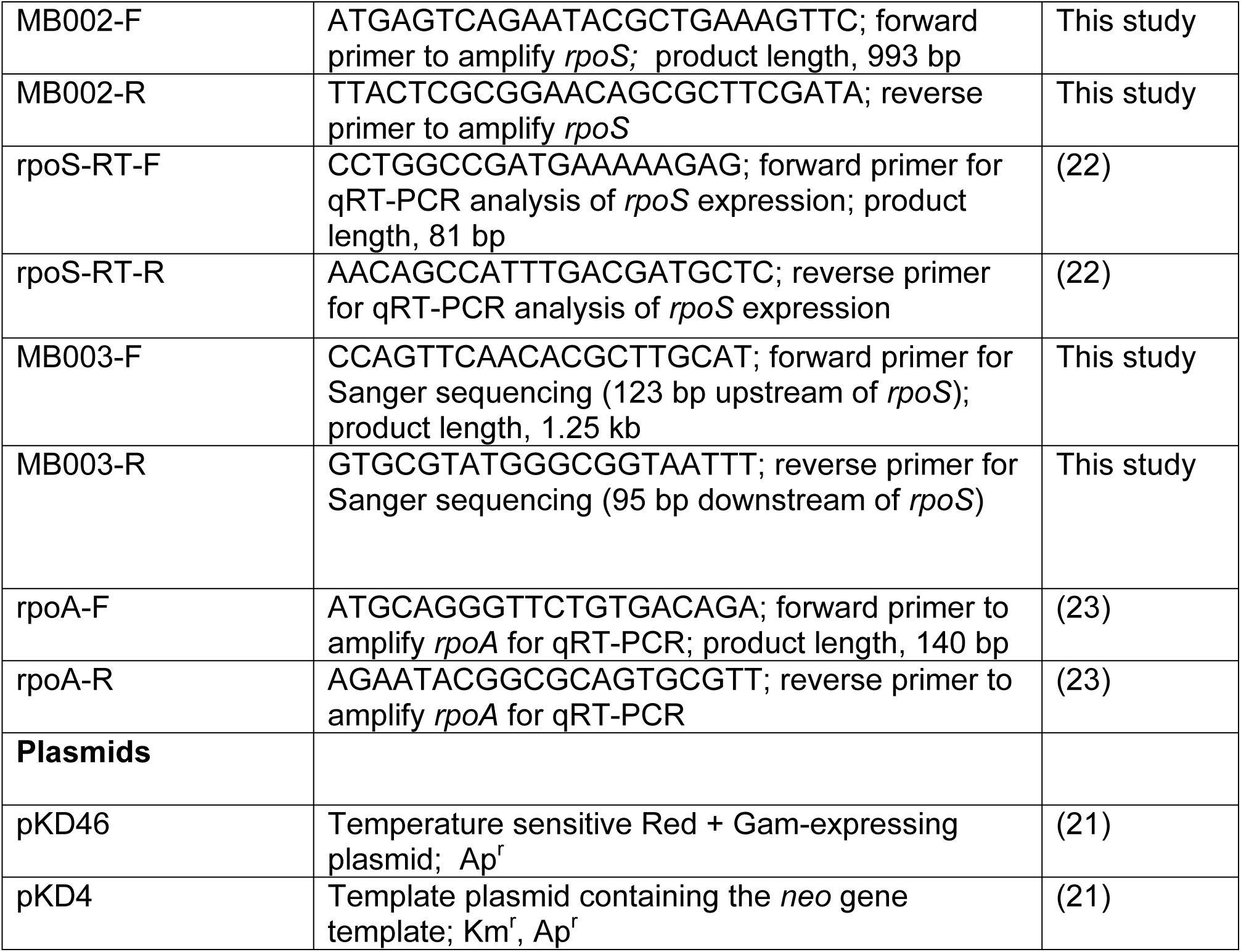
Bacterial strains, primers, and plasmids used in this study

### Real-Time Quantitative Reverse Transcription PCR (qRT-PCR)

qRT-PCR was used to verify loss of *rpoS* expression in the ∆*rpoS* ECOR2 mutant. Overnight cultures of wild-type and ∆*rpoS* ECOR2 were diluted 1:100 in sterile LB broth. After 8 hours of incubation, 1 mL of each culture was added to 2 mL RNAprotect Bacteria Reagent (Qiagen) according to manufacturer’s instructions. This timepoint was chosen because it correlates with entry into stationary phase of growth (Figure 1C and 1D), when RpoS is most abundant in the cell (19, 24). Samples were stored at -20°C until RNA extraction. RNA was extracted from each sample using the RNeasy Mini Kit (Qiagen) and quantified with the Quant-IT RiboGreen RNA Assay kit (Invitrogen). 1 ug of RNA was reverse transcribed to cDNA with the QuantiTect reverse transcription kit (Qiagen) according to the manufacturer’s instructions. For qRT-PCR analyses, 20 uL reactions were prepared using the QuantiTect SYBR Green PCR Kit (Qiagen) and primers rpoS-RT-F and rpoS-RT-R. qRT-PCR was performed on a LightCycler96 qPCR machine (Roche) with 45 cycles of 94°C for 15 seconds, 53°C for 30 seconds and 72°C for 30 seconds. Relative expression of *rpoS* was determined via the ∆∆Ct method using *rpoA* as the control gene (amplified with rpoA-F and rpoA-R). All reactions were followed by a melting curve to determine amplicon purity.

**Figure 1.**
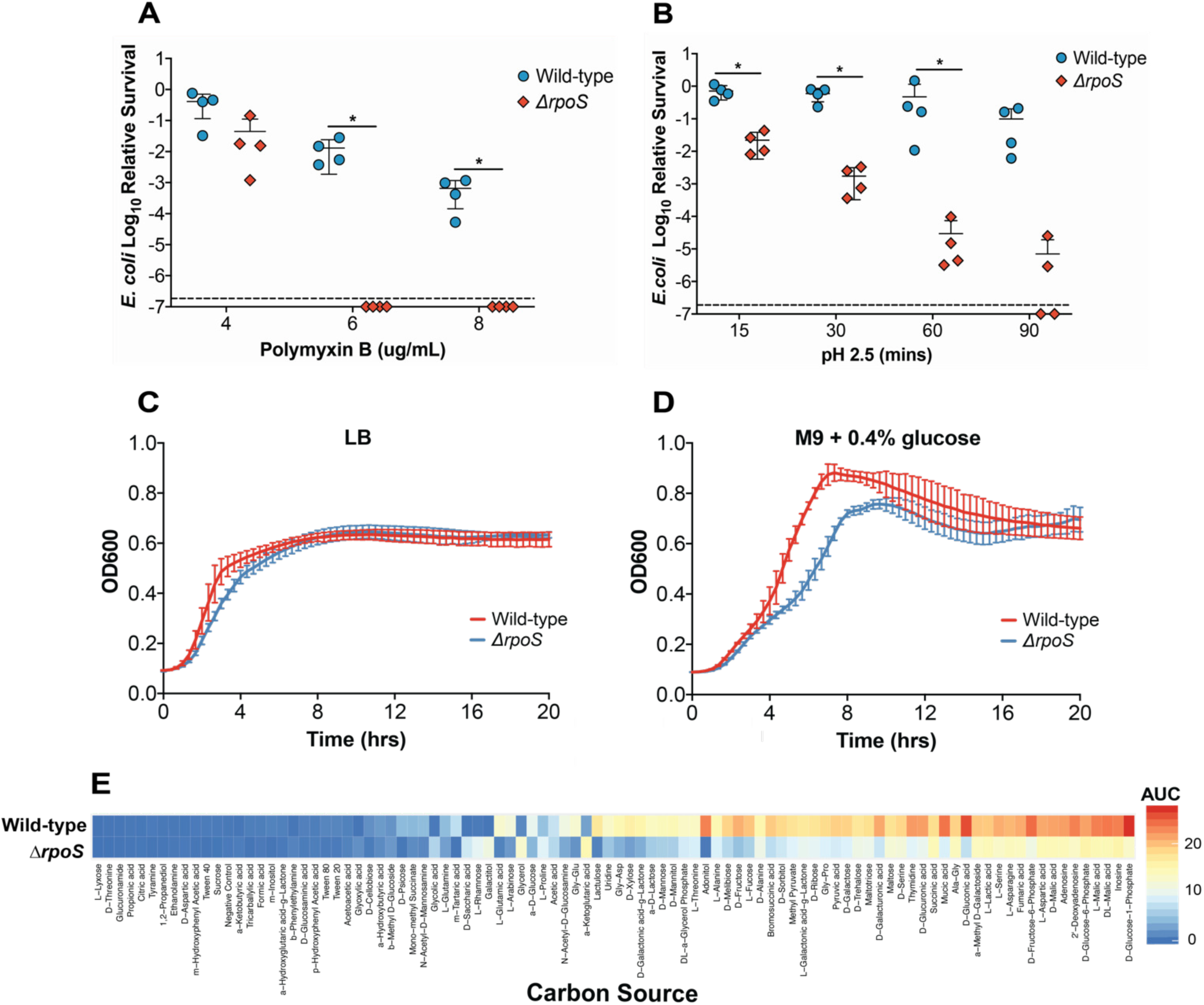
An isogenic ∆rpoS mutant of E.coli strain ECOR2 exhibits altered carbon metabolism and increased sensitivity to low pH, membrane stress, and nutrient limitation in vitro. (**A-B**) Wild-type ECOR2 and an isogenic *∆rpoS* mutant were tested for survival after one hour of treatment with 4, 6, and 8 ug/mL polymyxin B and after exposure to LB adjusted to pH 2.5 for 15, 30, and 60 mins. Relative survival was calculated by dividing the CFU from treated samples by that of the untreated control for each *E. coli* strain. The dashed line indicates limit of detection. N= 4 biological replicates per *E. coli* strain for each assay. Error bars represent standard deviation. p*<0.05, Mann Whitney U test or Welch’s unpaired t-test depending on data distribution. (**C-D**) Growth curves of wild-type ECOR2 and the *∆rpoS* mutant in LB or M9 supplemented with 0.4% glucose were conducted at 37ºC in a plate reader with shaking for 20 hours. Error bars denote standard deviation of the means of at least 3 biological replicates per *E. coli* strain. (E) Growth of wild-type ECOR2 and the *∆rpoS* mutant with 95 individual carbon sources using BiologTM PM1 plates. The metabolic capacity of each strain, depicted as a heat map, is represented as area under the curve (AUC) of OD595 values plotted over 24 hours for each carbon source.

### Phenotypic analyses

For growth curve analyses, overnight cultures of wild-type ECOR2 and the isogenic ∆*rpoS* mutant were diluted to OD_600_=0.01 in either sterile LB broth or M9 minimal media containing 0.4% glucose. 200 uL of each culture was transferred into a clear, flat-bottomed 96-well microplate. At least six technical replicates were included per experiment. A VersaMax microplate reader (Molecular Devices, LLC, Sunnyvale, CA) was used to measure OD_600_ at 20 minute intervals in microplates maintained at 37°C with regular shaking over a 20 hour time course.

To assess the ability of the ∆*rpoS* mutant to withstand stress *in vitro*, overnight cultures of the ∆*rpoS* mutant and its wild-type parent strain were diluted 1:100 in sterile LB broth and grown to mid-log phase (OD_600_ ~ 0.5-0.6). For membrane stress experiments, 500 uL of each culture was then treated with either 4, 6, or 8 ug/mL polymyxin B or water (as a control) and incubated for 1 hour. Polymyxin B binds lipopolysaccharide to destabilize the membrane of gram-negative bacteria (25). For pH stress assays, bacterial cultures were centrifuged at 4,000 xg for 5 mins, then 1 mL of cells was centrifuged a second time and re-suspended in LB Broth with a pH of 2.5 (adjusted with HCl). Cells were then incubated without shaking at 37°C for 15, 30, 60, and 90 minutes. Surviving cells were enumerated by serial plating on LB agar.

The metabolic properties of wild-type and ∆*rpoS* mutant ECOR2 was assessed with Biolog^TM^ PM1 (Biolog, CA) plates according to manufacturer’s instructions. Bacteria grown overnight on LB agar were resuspended in Biolog^TM^ inoculating fluid to a final OD_600_ of 1.0. Each well of the Biolog^TM^ plate was inoculated with 100 uL of cell suspension and incubated in a BioTek Kinetic plate reader (BioTek Instruments, Winooski, VT) for 24 hours. OD_595_ was used to measure reduction of tetrazolium violet dye every 15 minutes. Area under the curve (AUC) of OD_595_ values plotted over time were calculated as a measure of bacterial oxidation for each carbon source tested.

### HIO Generation and Experimentation

HIOs (13, 16) were generated as described previously (14) and maintained in media containing EGF, Noggin, and R-spondin (ENR media, see Ref. (15)) in Matrigel (8 mg/ml) without antibiotics prior to microinjection experiments. Bacterial cultures for microinjection were prepared by incubating wild-type ECOR2 and the ∆*rpoS* mutant overnight at 30°C with low shaking. The following day, cultures had reached an OD_600_ of ~1.0 and were diluted 1:10 in sterile PBS and centrifuged for 10 mins at 4000 xg. Bacterial cells were then re-suspended in sterile PBS or M9 minimal media supplemented with 1% galactitol, where indicated. 190 *μ*L of this suspension was mixed with 10 *μ*L of 4 kDa FITC-dextran suspended in PBS (2 mg/mL). FITC-dextran acted as a marker to ensure HIOs were successfully microinjected and the epithelial barrier remained intact (see below).

To introduce bacteria into the lumen of HIOs, microinjection was performed using thin-walled glass capillaries mounted on a Xenoworks micropipette holder with analog tubing, as previously described (Hill et al., 2017a, 2017b). Each HIO was microinjected with approximately 10^4^ CFU of wild-type ECOR2 or the isogenic ∆*rpoS* mutant. After microinjection, HIOs were incubated for 1 hour at 37°C. To remove bacteria introduced to the culture media during the microinjection process, HIO culture media was removed and cultures were rinsed with PBS and washed with ENR media containing 15 ug/mL gentamicin. HIOs were then washed again in PBS and the media was replaced with fresh antibiotic-free ENR. Successful microinjection was verified by visualizing fluorescence of FITC-dextran in each HIO several hours post-microinjection using an Olympus IX71 epifluorescent microscope. HIOs were considered successfully microinjected if a FITC signal was detectable. An absence in fluorescence was indicative of a loss in HIO structural integrity and these organoids were excluded from further analyses.

Bacteria were enumerated in the HIO lumen as previously described (26). Briefly, HIO culture media was removed from wells and each organoid was placed into individual screw-cap tubes containing 300 *μ*L PBS and 1.0 mm Biospec Zirconia/Silica beads (Fisher Scientific). HIOs were homogenized for 30 seconds in a Mini Bead Beater 8 (Biospec Products). Viable bacteria were enumerated via serial plating of the homogenate on LB agar with appropriate antibiotics.

### Germ-free mouse colonization

All animal experiments were performed with approval from the University Committee on Use and Care of Animals at the University of Michigan. Groups within an experiment were age-matched to the greatest extent possible. Male and female germ-free Swiss Webster mice aged 6-9 weeks old were obtained from a colony established and maintained by the University of Michigan Germ-free Mouse Facility. Mice received sterile food, water, and bedding and remained bacteriologically sterile (except for the experimental *E. coli* strains) throughout the course of the experiments. Bacterial inoculums were prepared as follows: Wild-type ECOR2 and the ∆*rpoS* mutant were grown in LB broth overnight at 30°C with low shaking. By the next morning, cultures had reached an OD_600_ of approximately 1.0. Cultures were then diluted 1:10 in sterile PBS and centrifuged at 4000 xg for 10 minutes. The supernatant was discarded and cells were resuspended in sterile PBS. Mice were inoculated via oral gavage with 100 *μ*L of bacterial suspension which equated to about 10^6^-10^7^ CFU per mouse of each individual strain for mono-colonization experiments and, for competition experiments, 10^6^ CFU per mouse of each strain administered together or sequentially (i.e. 24 hours apart), where indicated. Bacterial growth was measured by collecting and serial plating feces on LB agar. Feces were collected every other day for 7 days (mono-association studies) or 14-15 days (competition experiments).

### Statistical analyses

Area under the curve (AUC) values for Biolog assays and the corresponding heatmap were generated using custom R scripts (https://github.com/barronmr/Biolog_AUC.git). All other analyses were performed using GraphPad Prism 8.3 (GraphPad Software, Inc.). A Welch’s unpaired t-test was used for normally distributed data and a Mann Whitney U test was used for data that was not normally distributed. A P value of ≤ 0.05 was considered significant. Adobe Illustrator CC 2020 was used to arrange panels and generate final figures.

## Results

### Generation and *in vitro* characterization of an isogenic ∆*rpoS* mutant of *E. coli* strain ECOR2

The RpoS-mediated stress response has been well-studied in *E. coli* (19). Here, we generated an isogenic ∆*rpoS* mutant of *E. coli* strain ECOR2 (Figure S1A and S1B) to follow-up on our previous work assessing HIO epithelial responses to colonization by this strain (14). To verify the phenotype of the ∆*rpoS* mutant, we subjected it to various *in vitro* stressors. RpoS is known to protect against exposure to low pH (19), thus we first tested the ability of the ∆*rpoS* mutant to survive incubation in LB broth adjusted to pH 2.5. As expected, the *∆rpoS* mutant exhibited increased sensitivity to acid stress relative to the wild-type parent strain (Figure 1A). Incubation for one hour in LB broth containing 4, 6, or 8 ug/mL of polymyxin B, an antibiotic that binds lipopolysaccharide to destabilize bacterial membrane integrity (25), also revealed that the ∆*rpoS* mutant displayed a reduced ability to tolerate membrane stress compared to wild-type bacteria (Figure 1B). In addition, although it grew similar to wild-type ECOR2 under nutrient-rich conditions (Figure 1C), the ∆*rpoS* mutant exhibited a growth defect in minimal media supplemented with glucose as a carbon source (Figure 1D). To this point, a catabolic screen using Biolog^TM^ PM1 (carbon nutrition) microplates illustrated that the ∆*rpoS* mutant displayed a limited nutritional repertoire relative to wild-type ECOR2 (Figure 1E). These data support that loss of RpoS alters ECOR2 carbon metabolism and hinders growth during nutrient limitation. Together, the results from these experiments confirmed the importance RpoS for *E. coli* strain ECOR2 to withstand stressful environmental conditions.

### The HIO lumen is more restrictive to colonization by ∆*rpoS* ECOR2 compared to the germ-free mouse gut

The luminal microenvironment of HIOs is altered following microbial colonization, including a reduction in luminal oxygen concentrations and the deployment of epithelial antimicrobial defense mechanisms (14). To determine whether these changes pose stress to bacteria within the HIO lumen, wild-type ECOR2 or the isogenic ∆*rpoS* mutant was microinjected into individual HIOs and bacteria were enumerated after 24 hours. We found that the ∆*rpoS* mutant exhibited a significant defect in its ability to establish populations within HIOs (Figure 2).

**Figure 2.**
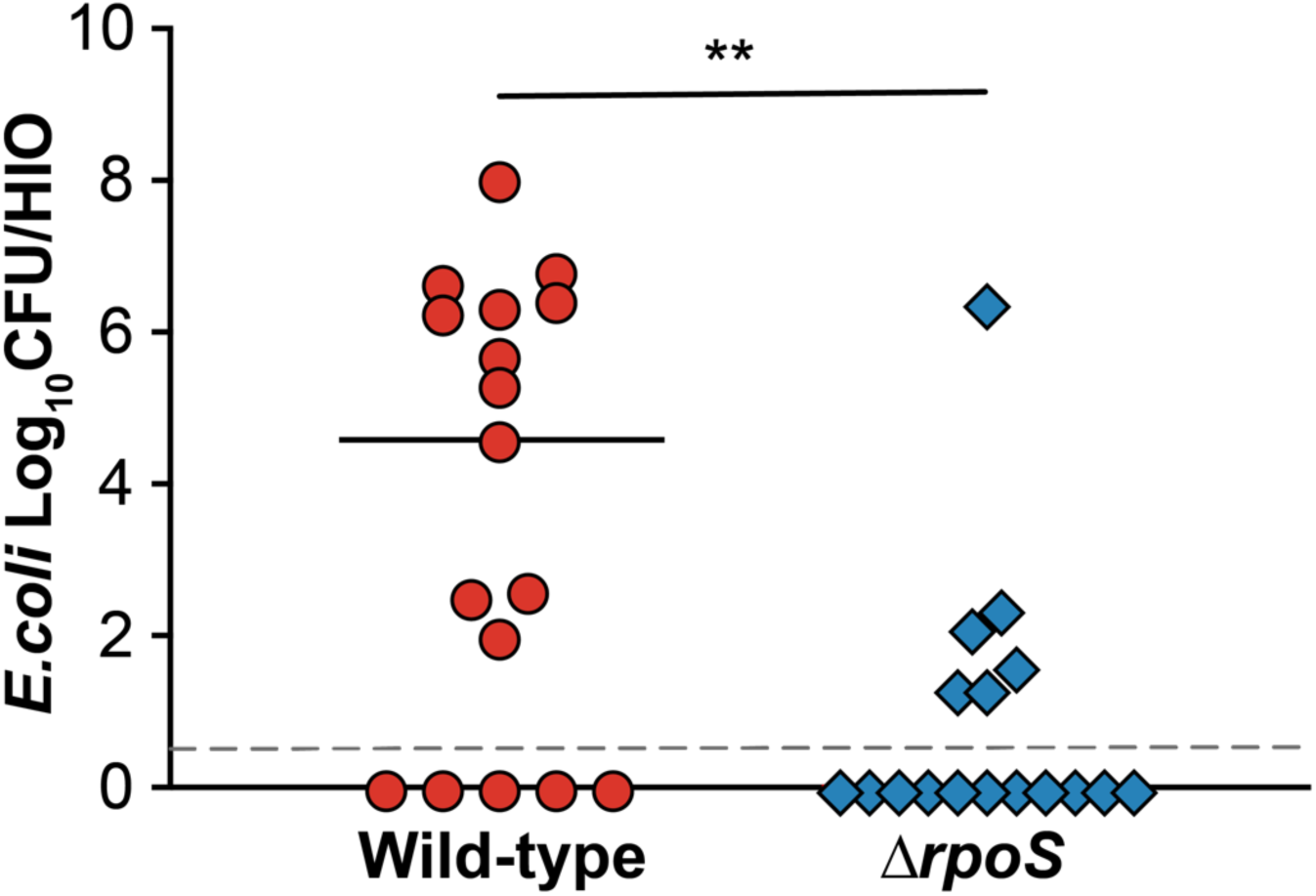
Loss of RpoS attenuates the ability of ECOR2 to colonize HIOs. Luminal CFUs obtained from individual HIOs 24-hours post-microinjection with ~104 CFU of either wild-type ECOR2 or its isogenic *∆rpoS* mutant. N= 17 biological replicates per *E. coli* strain. Error bars denote median; dashed line indicates limit of detection. **p<0.01, Mann-Whitney U test.

The stark colonization defect exhibited by the ∆*rpoS* mutant in HIOs prompted us to assess its colonization dynamics in a corresponding murine model. To do this, we colonized germ-free Swiss Webster mice with ~10^7^ CFU of wild-type ECOR2 or the *∆rpoS* mutant (Figure 3A). Feces were collected and plated every other day for seven days. Interestingly, in contrast to what we had observed in HIOs, the *∆rpoS* mutant colonized just as well as wild-type ECOR2 during mono-association of germ-free mice.

**Figure 3.**
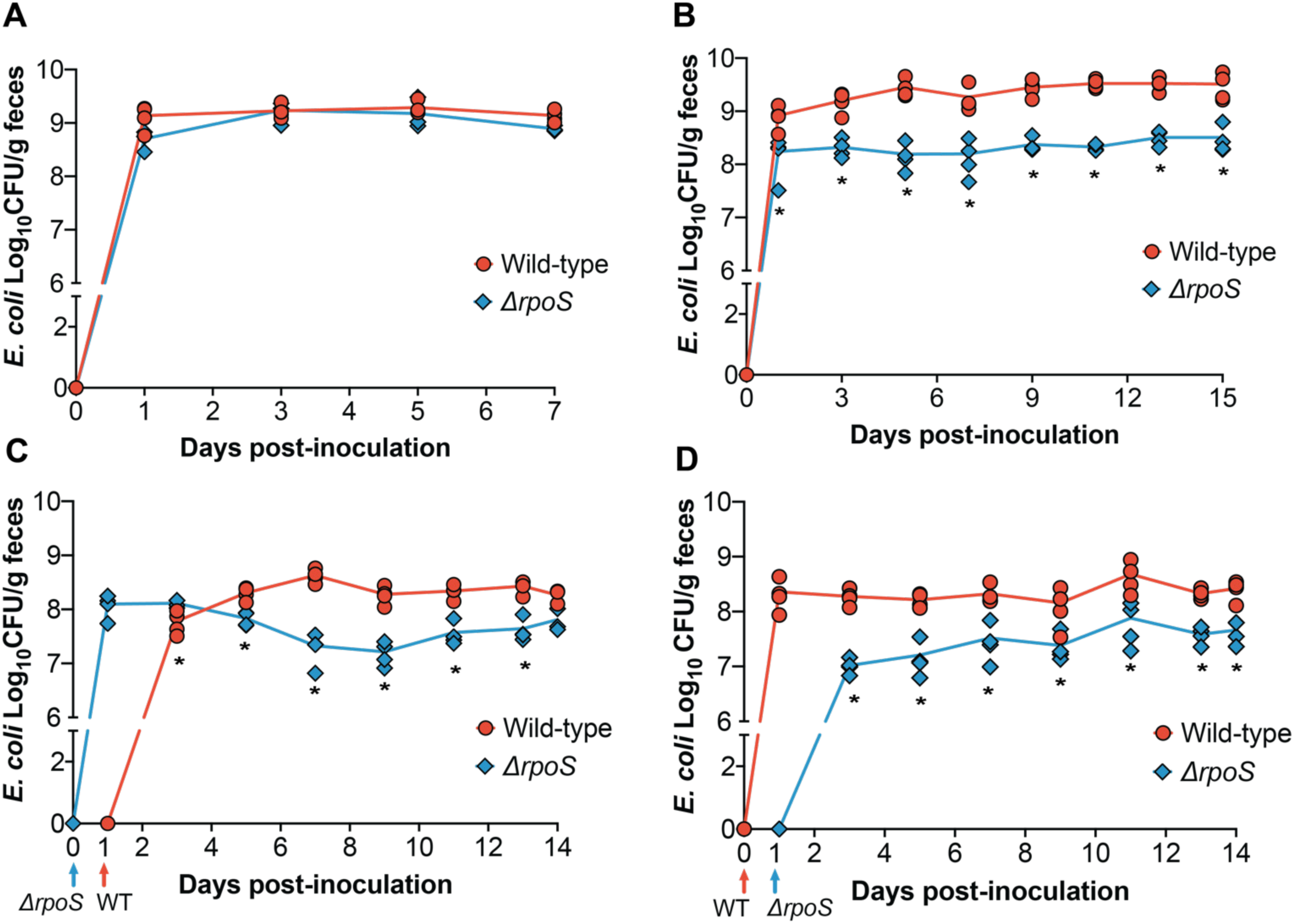
∆rpoS ECOR2 exhibits a colonization defect in the germ-free mouse gut only in the context of microbial competition. Germ-free Swiss Webster mice were either (**A**) mono-associated with 106-107 total CFU of wild-type or *∆rpoS* mutant ECOR2 or (**B**) competitively inoculated with a 1:1 ratio of each strain together or 24 hours apart, as demonstrated in (**C**) and (**D**). At the indicated times, fecal samples were homogenized, diluted, and plated as described in Materials and Methods. N= 3-4 mice per timepoint. *p<0.05, Mann-Whitney U tests.

We next asked whether the colonization dynamics of the *∆rpoS* mutant would change during competition with the wild-type strain. We posited that the relative stress within the gut environment would increase in the context of microbial competition, and thus could influence the ability of *∆rpoS* to stably colonize the intestine. To address this question, we colonized germ-free mice with equal amounts of wild-type and ∆*rpoS* ECOR2 (total CFU ~ 10^6^) and measured fecal levels of each strain over fifteen days. While the ∆*rpoS* mutant was still able to colonize the gut, it exhibited a fitness defect during co-colonization with wild-type ECOR2 (Figure 3B). We hypothesized that allowing the *∆rpoS* mutant to establish within the germ-free mouse gut prior to introduction of the wild-type strain would rescue this defect. To test this hypothesis, mice were colonized with *∆rpoS* followed by the wild-type strain 24 hours later, or vice versa (Figure 3D and 3E). Feces were collected and plated over fourteen days. We found that the *∆rpoS* mutant displayed a competitive disadvantage regardless of whether it was introduced before (Figure 3D) or after (Figure 3E) the wild-type strain, thus indicating that order of colonization does not matter.

## Discussion

In this study, we used an isogenic *∆rpoS* mutant of *E. coli* strain ECOR2 as a biological probe to investigate the bacterial response to colonization of the HIO lumen relative to the germ-free mouse gut. We chose to focus on *E. coli* for several reasons. First, we previously used ECOR2 to study the response of the HIO epithelium to initial colonization (14) and we wished to examine the microbial side of this interaction. Second, and more importantly, multiple studies have demonstrated that *E. coli* is one of the first bacterial species to establish within the gut after birth (2, 27, 28). Thus, *E. coli* represents an appropriate organism to study in the context of the microbe-naïve environment modelled by HIOs and germ-free mice.

In *E. coli* and other Gram-negative bacteria, the ability to respond to diverse stressors in the environment occurs via the RpoS-mediated stress response (18–20). Indeed, our data demonstrate that loss of RpoS decreases the ability of *E. coli* strain ECOR2 to withstand nutrient limitation, membrane stress, and exposure to low pH. While RpoS has been extensively studied in the context of laboratory-imposed environmental stress, it has also been shown to facilitate bacterial colonization of the gut. For example, RpoS is important for efficient intestinal colonization by several pathogenic bacterial species, including *Vibrio cholerae*, *Salmonella enterica* serovar *typhimurium,* and the virulent *E. coli* strain O157:H7 (29, 30). In addition, it was previously reported that RpoS may play a role in the initial establishment of non-pathogenic *E. coli* within the streptomycin-treated mouse gut (31). Thus, the ability to mount a global, cross-protective stress response can be advantageous during colonization of the intestine, where microbes must sense and respond to conditions ranging from nutrient limitation to host antimicrobial defense mechanisms. Given the role of RpoS in colonization of the intestine, we leveraged this information to investigate the relative stress imposed by the HIO lumen on colonizing bacteria. We found that loss of RpoS significantly decreased the ability of ECOR2 to colonize HIOs, though it did not prevent colonization of germ-free mice. These data indicate that the type or severity of microbial stressors within HIOs differ from those of the *in vivo* intestinal environment.

There are several possible explanations for the colonization dynamics exhibited by the *∆rpoS* mutant in HIOs versus germ-free mice. Our Biolog^TM^ screen revealed that the *∆rpoS* mutant had a more limited nutritional repertoire relative to wild-type ECOR2. With this in mind, one hypothesis is that the HIO lumen is more restricted in terms of nutrient availability compared to the murine intestine. To this end, the mouse gastrointestinal tract contains a relatively rich nutrient pool that is replenished during regular feeding (32,33). The nutrient milieu of HIOs is relatively unknown, though it is presumably derived from the surrounding culture media that has been further modified by the metabolic activities of the epithelium (34) and mesenchyme. Thus, the diversity and concentration of substrates available for *E. coli* consumption within HIOs may be limited. Preliminary evidence supporting this hypothesis comes from our observation that microinjecting the *∆rpoS* mutant into HIOs in minimal medium supplemented with galactitol, a nutrient identified through our Biolog^TM^ assays that supports *∆rpoS* mutant growth, partially rescues its colonization defect relative to bacteria microinjected in PBS (Figure S1C).Therefore, providing *E. coli* a nutritional advantage may help alleviate at least one source of stress within HIOs. However, more extensive work is required to determine the specific substrates capable of mitigating the potential nutritional deficits of the HIO lumen.

Nutrient availability may also explain the colonization dynamics observed *in vivo,* where the *∆rpoS* mutant grew to wild-type levels during mono-association of germ-free mice yet displayed a colonization defect in the context of microbial competition. Indeed, while a number of factors contribute to bacterial intestinal colonization, the ability to compete for a limited number of nutritional niches is paramount for successful colonization of the gut (35,36). In addition, several studies have demonstrated that nutrient availability is the primary driver of *E. coli* colonization and adaptation within the murine gut (9, 35, 37). Thus, competition may increase the nutritional stress within the germ-free mouse intestine, though perhaps not to the same degree as that inherent to the HIO lumen. Indeed, the fact that the *∆rpoS* mutant was able to establish relatively robust populations in *in vivo* suggests that, in contrast to HIOs, the nutritional milieu of the germ-free mouse gut is diverse enough to support bacteria with different metabolic flexibilities and efficiencies. Comparing the ability of *E. coli* mutants deficient in specific metabolic pathways to colonize HIOs and germ-free mice would provide a more targeted understanding of the nutritional stressors present, or absent, within each system.

Another possible explanation for our results is that the lack of peristalsis and luminal flow within HIOs, which leads to a build-up of epithelial and microbe-derived waste within the HIO lumen (38), creates a relatively stressful environment for colonizing microbes. To this point, a number of technological advancements have been made with regard to the stationary nature of the HIO system. For example, several studies have demonstrated that organoids can be dissociated and seeded onto microfluidic devices, or “chips”, that allow for controlled and continuous media flow across the resulting cell monolayer (39–41). While these intestine-on-a-chip technologies mimic conditions of the *in vivo* gut, they lack the 3-D HIO structure that can be advantageous for certain applications, including microbial colonization experiments. To this end, Sidar et al. recently developed a multifluidic device that “ports” HIOs to establish steady-state liquid flow through the lumen for several days, thus eliminating luminal waste while maintaining the structural integrity of 3-D organoids (38). These examples illustrate how HIOs can be manipulated to better recapitulate *in vivo* intestinal structure and physiology, thus creating novel tools for investigating mechanisms modulating epithelial-microbe interactions within the gut (12). In this regard, our *∆rpoS E. coli* mutant, or other bacterial “probes,” can serve as a means of comparing these emerging systems and provide insight into how closely they mirror the *in vivo* gut environment from a microbial perspective.

Overall we have demonstrated that, from a bacterial standpoint, HIOs possess a unique luminal environment relative to the germ-free mouse intestine. Our results indicate that the type or severity of microbe-perceived stress within the HIO lumen differs from that of the *in vivo* gut environment. While we did not identify the specific conditions responsible for these differences, we suspect that there are a number of factors that collectively shape the distinct environment of the HIO lumen. Ultimately, the results from this study better characterize the HIO model system as it pertains to studying the establishment of bacterial populations at the intestinal epithelial interface. Moving forward, such knowledge will help inform when and how we use HIOs to investigate diverse aspects of host-microbe symbioses within the gut.

## Acknowledgements

We acknowledge the University of Michigan Germ-free Mouse Core for assistance with mouse colonization experiments. We also thank Stephanie Thiede and Emily Benedict for providing R scripts for Biolog^TM^ assay analyses, as well as the laboratories of Mary O’Riordan and Harry Mobley for sharing bacterial strains and equipment. Finally, we thank Kimberly Vendrov, Annie Pinchkoff, Michelle Smith, and Pariyamon Thaprawat for experimental assistance.

This work was funded by the National Institutes of Health cooperative agreement AI116482 to V.B.Y/J.R.S. and AI007528 to M.R.B.

The funders had no role in study design or data collection and interpretation.

VBY has served as a consultant to Vedanta Biosciences, Bio-K+ International and Pantheryx.

## Author contributions

M.R.B, R.J.C, and V.B.Y, conception or design of work; M.R.B., R.J.C., D.R.H., S.H., V.K.Y, data collection; M.R.B. and R.J.C, data analysis; M.R.B., V.B.Y drafting the article; M.R.B., R.J.C., D.R.H., S.H., V.K.Y., J.R.S., and V.B.Y. critical revision of manuscript; M.R.B., R.J.C., D.R.H., S.H., V.K.Y., J.R.S., and V.B.Y., final approval of version to be published.

